# Shape-Dependent Toxicity of Gold Nanoparticles in Microalgae: Distinct Cellular and Molecular Responses

**DOI:** 10.64898/2025.12.15.694526

**Authors:** Can Wang, Bhaskar Sharma, Ruonan Peng, Xin Yong, Louis S. Santiago, Ke Du

**Affiliations:** Department of Chemical and Environmental Engineering, University of California Riverside, CA, 92521, United States; Department of Mechanical and Aerospace Engineering, University at Buffalo, NY, 14260, United States; Department of Botany and Plant Sciences, University of California Riverside, CA, 92521, United States

**Keywords:** Nanoparticles, Morphology, Microalgae, Ecotoxicity, Cellular and molecular mechanism, Transcriptomics

## Abstract

The ecotoxicological effects of engineered nanoparticles in aquatic environments are influenced not only by composition and size but also by particle morphology, yet shape dependent interactions with primary producers, remain poorly understood. In this study, we evaluated the cellular and molecular responses of freshwater algae (*Chlamydomonas reinhardtii*) following exposure to 100 nm silica-coated gold nanospheres (AuNP) and nanostars (AuNS) across multiple concentrations. Exposure to 10 mg/L AuNS for 72 h results in significantly stronger inhibition of algal growth and photosynthetic activity compared to the same concentration of AuNP. Morphometric profiling reveals that AuNS induced pronounced structural injury, including cell enlargement, debris production, and disruption of subcellular organization than AuNP. Confocal imaging suggested this heightened toxicity may stem from distinct internalization patterns, with AuNP primarily adhering to chloroplast surfaces, whereas AuNS penetrated more deeply into intracellular compartments. RNA sequencing identified 9 upregulated and 38 downregulated differentially expressed genes (DEGs) in the 10 mg/L AuNP treated cells, impairing photosynthesis and energy storage via the photosystem II subunit S1 (PSBS1)/ early light-inducible protein (ELI3) pathway. In contrast, the AuNS group exhibits 246 upregulated and 145 downregulated DEGs, affecting membrane integrity and nitrogen metabolism through the nitrate reductase (NIT1)/ aminomethyl transferase (AMT1)/ protein kinase domain-containing protein (A0A2K3CRU5) pathway. These results demonstrate that nanoparticle morphology can drive divergent toxicity mechanisms in algal cells. Our findings highlight the necessity of incorporating NPs morphology into environmental risk assessments and suggest that safer nanomaterial design should consider shape-dependent interactions with aquatic microorganisms.

**Graphic Abstract:** 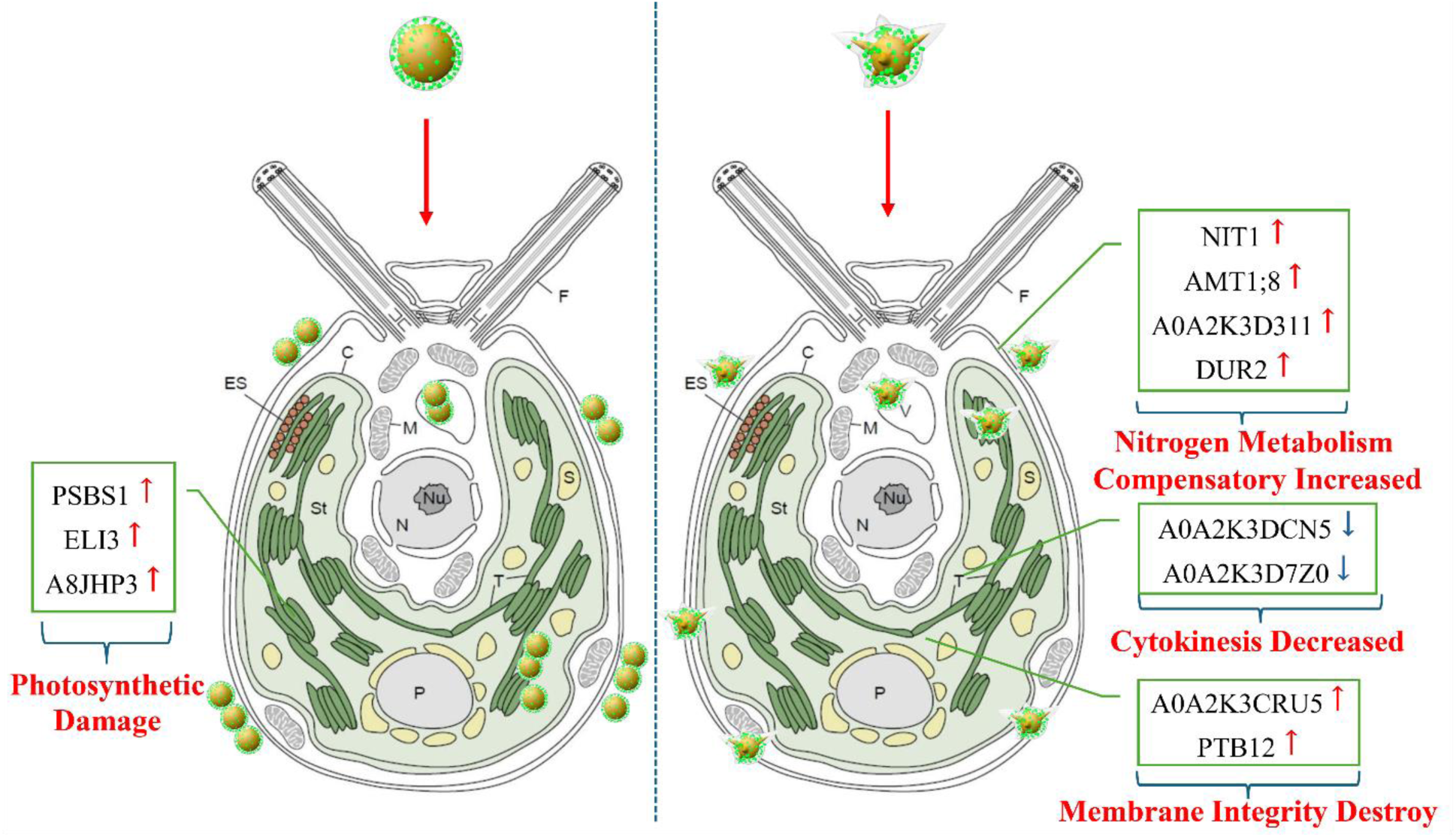

## Introduction

Nanoparticles (NPs) are tiny materials with size ranges from 1 to 100 nm^1^. Their high surface area and nanoscale dimensions confer unique physical and chemical properties, making them widely utilized in a variety of commercial and research applications, such as catalysis^2^, imaging^3^, medical applications^4^, energy-based research^5^, and environmental applications^6^. The distinctive morphological characteristics of NPs have attained great interest since morphology influences most of the physicochemical properties^7^. For instance, the morphology of NPs plays a critical role in modulating the rate and mechanism of cellular uptake and the intracellular transport^8, 9^, and the shape of NPs also influences the drug delivery^10^, cell-nanoparticle interaction and internalization rate^11^, and even the behavior or differentiation of stem cells^12^. These properties prompted research on precisely controlling the shape of NPs to understand and customize these nanostructures for desired application^13^.

In the process of NPs production, usage, transportation, and disposal, they inevitably enter the environments through migration and transformation^14^. The aquatic environment, which receives runoff and wastewater from domestic, medical, and industrial sources, is the primary medium through which NPs enter and diffuse into other environmental resources^15^. Current estimates indicate that 66,000 metric tons of NPs are released into surface waters annually^16^. Numerous studies mentioned that the shape of NPs influences systemic toxicity, biodistribution and biological activity, manifesting in shape-induced directed differentiation^17^, cellular death via apoptosis^18^, necrosis^19^, gene transfection and transfer^20^, and metabolism alteration^21^. These effects arise from different surface areas, uptake level, protein corona, physical disruption of the cell membrane, and wettability and surface curvature of the particles^22^. Consequently, the wide usage and massive discharge of NPs with different shapes makes it imperative to determine their toxic effects in aquatic environment. However, there remains limited information regarding the ecological hazards posed by NPs of different shapes.

The World Health Organization (WHO) highlighted the concept of ecological public health^23^, calling attention to the threats to human health caused by global ecological environment changes, such as climate change, biodiversity loss, alterations in hydrological systems, freshwater depletion, and contamination^24^. Microalgae play a crucial role in mitigating these eco-environmental changes. They contribute significantly to regulating climate change by producing up to 60% of the world’s oxygen through photosynthesis and controlling atmospheric carbon dioxide levels^25^. Additionally, as the foundation of the aquatic food chain, the abundance and quality of microalgae are essential for maintaining biodiversity. Moreover, the health of microalgae directly influences water quality and ecological resilience^26^. *Chlamydomonas reinhardtii (C. reinhardtii)*, a model organism extensively studied due to its widespread distribution in soil and freshwater, was selected for this study to investigate the varying impacts of NPs with different shapes on the ecological environment.

In recent decades, gold nanoparticles (AuNPs) of different shapes have been used widely for specific biomedical applications due to their distinct optical properties, such as drug delivery^27^, gene therapy^28^, diagnose of diseases^29^, and bio-imaging^30^. Many studies have demonstrated that the toxicity of AuNPs toward microalgae is multifactorial, influenced by particle size, surface coating, concentration, exposure time, and morphology. For instance, exposure of *C. reinhardtii* to 42 nm AuNPs at 5 mg/L for 24 hours results in slight growth inhibition, whereas a concentration of 20 mg/L induces oxidative stress in approximately 15% of cells^31^. *Raphidocelis subcapitata* exposed to 30 nm AuNPs for 72 hours exhibits an EC_50_ of approximately 350 mg/L^32^. EC_50_ value after 30 min of treatment was 114 mg/L Polyamidoamine-coated AuNPs for *C. reinhardtii*^33^.

Despite these insights, studies addressing the role of nanoparticle shape in algal toxicity remain limited. Silica is frequently employed as a stabilizing surface coating for AuNPs, as the silica shell can stabilize the core of AuNPs^34, 35^ by preventing non-specific interactions with biomolecules^36^. Both silica and gold are considered biocompatible, inert and nontoxic materials^37^, thereby minimizing intrinsic material toxicity to cells such as microalgae, and making silica-coated AuNPs (Si-AuNPs) ideal for studying shape-dependent algal toxicity.

In this study, 100 nm star (AuNS) and spherical (AuNP) Si-AuNPs were used as the representative shapes of NPs, and exposed *C. reinhardtii* at varying concentrations to investigate their differential effects on algal growth and primary photosynthetic processes. Confocal and transmission electron microscope were employed to detect the interaction and accumulation of different shapes of AuNPs and *C. reinhardtii.* Furthermore, RNA sequencing (RNA-seq) quantifies genome-wide gene expression, which is essential for interpreting the functional elements of the genome and understanding development and disease^38^. Therefore, RNA-seq was employed to elucidate the unique toxicity mechanisms of these differently shaped AuNPs in microalgae.

## Materials and Methods

### Nanoparticle characterization

Silica-coated gold nanospheres (AuNP) and gold nanostars (AuNS) (100 nm in size) were purchased from CD Bioparticles (NY, USA). Gold nanoparticles (AuNPs) were monodispersed in DI water to form a suspension with a concentration of 1, 10, and 50 mg/L for exposure experiments. The detailed production information and number of particles in each group are provided in Supplementary Materials Table S1. The AuNPs solution was ultrasonicated for at least 10 min before use. All AuNPs suspension samples were stored at 4 °C. Transmission electron microscopy (TEM) was employed to observe the size and shape of the AuNPs. Zeta potentials of AuNPs were measured by phase analysis light scattering (Malvern Zetasizer Nano ZS), using disposable cuvettes. Measured zeta potentials averaged over 3 runs, for each calibration step at least 300 particles were measured with typical particle concentrations of 10^9^–10^10^ particles/ml^39^. The surface plasmon resonance (SPR) peaks were measured using a spectrophotometer (BioTek, CA, USA).

### Algal cultivation and AuNPs exposure

Wild type *Chlamydomonas reinhardtii* (CC-125) was obtained from the Algae Research Supply Company (Carlsbad, CA, USA). The strain was activated in Tris-Acetate-Phosphate (TAP) medium. The strain was activated in Tris-Acetate-Phosphate (TAP) medium. *C. reinhardtii* were inoculated into fresh medium and grown at 25 ± 1 °C and an incident light intensity of 4800 lx light intensity at the surface of the cultivation vessel. The light-dark regime was set to 12 h of light and 12 h of darkness. Cell density was maintained by dilution to 1×10^6^ cells/ml at the beginning of each light cycle. Cultures were maintained under the same conditions until to the timepoint of experiments (i.e. 24, 48, 72 h) to obtain highly synchronous cultures^40^. When algal culture was in the exponential growth phase, an aliquot having a cell density of 1×10^6^ cells/ml was used for treatments. Cell density was measured using a spectrophotometer at a wavelength of 680 nm and represented by the optical density (OD 680) values. All flasks were shaken three times each day and their locations were randomly shifted. The algae were exposed to varying concentrations of AuNPs (1, 10, and 50 mg/L) or deionized water (vehicle control). Given that exposure to 42 nm AuNPs for 24 h induced oxidative stress of *C. reinhardtii*^31^, while exposure to 30 nm AuNPs for 72 h produced an EC_50_ of approximately 350 mg/L of *C. reinhardtii*^32^, we established 1 mg/L (≈ 1/100 EC_50_) as a low dosage, 10 mg/L (≈1/50 EC_50_) as a medium dose, and 50 mg/L (≈1/10 EC_50_) as a high dose. These concentrations were selected to generate an appropriate dose–response relationship and to capture a comprehensive range of toxicological effects across hierarchical exposure levels.

### Cell viability

Algal cell density was counted using a Blood Counting Chamber under microscopy. Cell growth rate was measured using a spectrophotometer at a wavelength of 680 nm at 0 h and 24 h after culturing in TPA medium and represented by the optical density (OD_680_) values. The growth rate was calculated using the following equation:

Growth rate (%) = (OD_24h_-OD_0h_)/OD_0h_ × 100%.

### Determination of photosynthetic pigments concentration

After centrifuging the *C. reinhardtii* suspension at 24 h, the algae pellet was resuspended in the same volume of 80% acetone. Then, the tube was heated in a water bath at 55 ℃ in the dark for 30 minutes. The absorbance values of supernatant were measured at 470, 645, and 663 nm. The formula for calculating photosynthetic pigments content is as follows^41^:

Chlorophyll a (C_a_) = 12.21 A_663_ – 2.81 A_645_

Chlorophyll b (C_b_) = 20.13 A_645_ – 5.03 A_663_

Carotenoids (C_c_) = (1000 A_470_ - 3.27×C_a_-104×C_b_)/229

### Measurement of Electron Transport Rate (ETR)

After dissolving microalgae into TAP medium for 24 h, we characterized photosynthesis with chlorophyll fluorescence-light curves^42^ using a photosynthesis yield analyzer (Mini-PAM, Heinz Walz GmbH, Effeltrich, Germany). We constructed chlorophyll fluorescence light response curves using photon flux density (PFD) values of 0, 55, 81, 122, 183, 262, 367, 616, and 1115 μmol m^-2^ s^-1^ to bring the light up to the highest level, and calculated maximum quantum yield of photosystem II (Y(II)) and electron transport rate (ETR) at the highest light level^43^. ETR was calculated based on chlorophyll fluorescence data using the equation: ETR = Y(II) × PFD × 0.84 × 0.5, where Y(II) is the effective quantum yield of photosystem II (PSII) measured during a 0.8-s saturating flash (6,000 μmol m^-2^ s^-1^), with an average ratio of light absorbed of 0.84 an assumed average ratio of photosystem II to photosystem I reaction centers of 0.50^44^.

### Flow cytometry

FC was performed in a device (MoFlo Astrios EQ, CA, USC). Before running in the flow cytometer, samples were filtered by 35 μm nylon mesh screen and shacked for 10 s to prevent aggregate formation and subsequent clogging of the laminar flow chamber. Samples were diluted adjusting the dilution factor to ensure a suitable number of events, 1,000 per second. Forward angle light scatter (FSC) and right-angle light scatter (SSC) were plotted in a logarithmic scale and data were analyzed with NovoExpress 1.5.6 software. Average FSC, SSC, and cell population percentages were calculated. The *C. reinhardtii* cells populations and debris were discriminated based on the FSC versus SSC plot.

### Confocal laser Scanning Microscopy (CLSM)

*C. reinhardtii* were co-cultured with AuNPs for 72 h and observed on a TCS-SP5 Spectral confocal laser scanning microscope (Leica Microsystems, Heidelberg, Germany), using the 63× objective(oil immersion) lens and excitation was performed via a 488 nm line of an argon ion laser and a 561 nm diode laser. For imaging, algal cells were mixed with 1% agarose (cooled to room temperature) and mounted on a glass slide. The optimized emission detection bandwidths were configured at 500–550 nm (AuNPs-AF488) and 680–780 nm (autofluorescence of chloroplasts) by a hybrid detector. Zstack section thickness was set to 0.4 to 0.6 μm. The captured images were processed through Fiji software package. The colocalization of AF488 labelled nanoparticles with chloroplast in confocal images was performed by using Coloc 2 function in Fiji2 software. The correlation between fluorescent signals was analyzed using Manders’ overlap coefficient.

### Transmission Electron Microscope (TEM)

TEM was used to observe the size and shape of the AuNPs and the morphology and subcellular structure of *C. reinhardtii* after exposure at 72 h. After a 72 h exposure to AuNPs, *C. reinhardtii* cells were harvested as cell pellets and subsequently fixed with 2.5% glutaraldehyde in 0.1 M sodium cacodylate buffer (SC buffer, pH 7.4). Samples were then treated with 1% osmium in 0.10 M SC buffer for 1 h on ice and washed in 0.10 M SC buffer. Ultrathin sections (60 nm) were cut using a Leica microtome with diamond knife and followed by post-staining with both uranyl acetate and lead. Images were captured on a JEOL 1400 plus TEM at 80 kV with a Gatan One view 4k × 4k camera.

### mRNA sequencing

We collected algae after 72 h exposure to 10 mg/L AuNP and AuNS for RNA-seq to analyze mRNA expression. Total RNA was isolated and purified using TRIzol reagent (Invitrogen, CA, USA) following the manufacturer’s procedure. Each sample’s RNA amount and purity was quantified using NanoDrop ND-1000 (NanoDrop, DE, USA), and the RNA integrity was assessed by Bioanalyzer 2100 (Agilent, CA, USA). Poly (A) RNA was purified from 50 μg total RNA using Dynabeads Oligo (dT) 25-61005 (Thermo Fisher, CA, USA) by two rounds of purification. Then, the poly(A) RNA was fragmented into small pieces using Magnesium RNA Fragmentation Module (NEB, cat. e6150, USA) under 86℃ for 7 min. An A-base was added to the blunt ends of each strand in preparation for the ligation of the indexed adapters. Dual-index adapters were ligated to the fragments, and size selection was performed with AMPureXP beads. After heat-labile UDG enzyme (NEB, cat. m0280, USA) treatment of the U-labeled second-stranded DNAs, the ligated products were amplified with PCR by the following conditions: initial denaturation at 95℃ for 3 min; 8 cycles of denaturation at 98℃ for 15 sec, annealing at 60℃ for 15 sec, and extension at 72℃ for 30 sec; and then final extension at 72℃ for 5 min. Finally, we performed paired-end sequencing (PE150) on an Illumina Novaseq™ 6000 platform (CA, USA) following the vendor’s recommended protocol, sequencing depth is 35-45 million mapped reads per sample. The sequence quality of the mRNA samples were verified using FastQC (https://www.bioinformatics.babraham.ac.uk/projects/fastqc/) and RseQC (http://rseqc.sourceforge.net/), Q30 scores >=93%). Sequence reads were trimmed to remove possible adapter sequences and nucleotides with poor quality using Trimmomatic v.0.36. The trimmed reads were mapped to the_*Chlamydomonas_reinhardtii* reference genome available on ENSEMBL using the STAR aligner v.2.5.2b. The STAR aligner is a splice aligner that detects splice junctions and incorporates them to help align the entire read sequences. BAM files were generated as a result of this step. Below are the statistics of mapping the reads to the reference genome. Unique gene hit counts were calculated by using featureCounts from the Subread package v.1.5.2. The hit counts were summarized and reported using the locus_tag feature in the annotation file. Only unique reads that fell within exon regions were counted. After extraction of gene hit counts, the gene hit counts table was used for downstream differential expression analysis. Using differential-expression (DEseq2) statistics, a comparison of gene expression between the different groups of samples was performed. The differentially expressed transcripts and genes were selected with log^2^ (fold change) ≥ 1 or log^2^ (fold change) ≤ -1 and *p* value < 0.05 criteria with the R package edgeR (https://bioconductor.org/packages/edgeR).

## Statistical Analysis

Statistical differences were performed using a Student’s t test for two-group comparisons or one-way ANOVA for multiple comparisons with a Student−Newman−Keuls test. Data are presented as mean ± SD, and *p* < 0.05 was considered statistically significant.

## Results

### Properties of AuNP and AuNS

In this study, two representative morphologies of gold nanoparticles were utilized: 100-nm-diameter silica coated gold nanospheres (AuNP), and 100-nm-diameter silica coated gold nanostars (AuNS). For fluorescence observation, fluorophores (Ex/Em: 488/514 nm) were attached to naked AuNP and AuNS with 80 nm (core) diameters. To ensure the stability of the AuNPs, a silica coating was applied, rendering a final particle size of 100 nm (Figure 1A). The Transmission electron microscopy (TEM) measured morphology of the particles was in agreement with the morphology reported by the supplier. TEM imaging confirmed successful functionalization and high homogeneity of the AuNPs (Figure 1B). Particle size distribution obtained from the TEM image analysis were fitted to the log-normal distribution. The peak values of the fit were taken as the average particle size of the corresponding samples. Two kinds of NP, as a result had similar size, at the range of 100 ± 14 nm (Figure 1C). The AuNP and AuNS displayed shape-dependent surface plasmon resonance (SPR) peaks at approximately 580 nm and 800 nm, respectively (Figure 1D). Both two kinds of particles exhibited a negative zeta potential as AuNP were −23.2 mv and −16.8 mv for AuNS.

**Figure 1.**
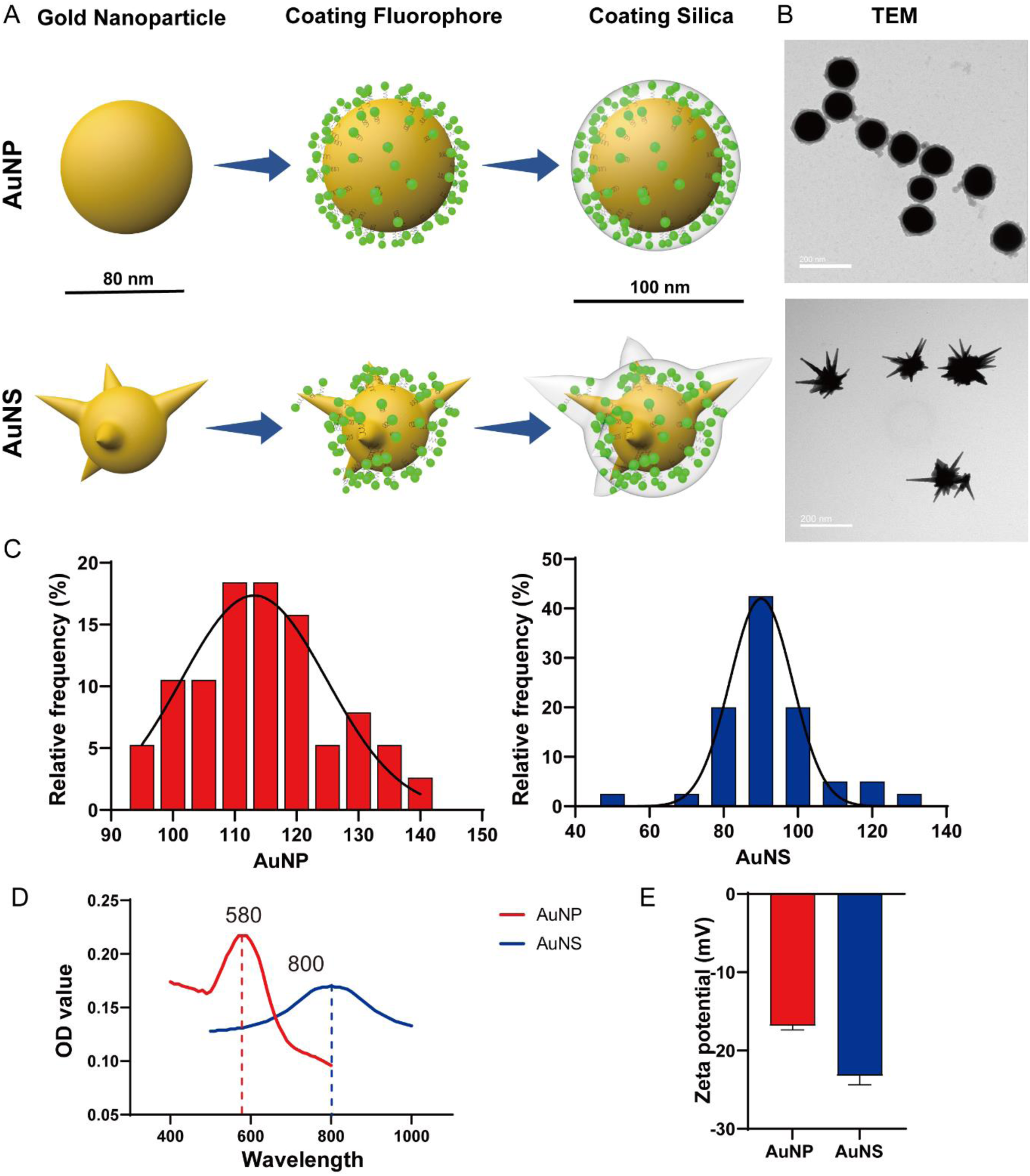
Silica coated-AuNPs preparation and characterization. A. AuNPs samples used in this study: 100-nm-diameter AuNP and AuNS. B. TEM characterization. The scale bar is 200 nm. C. Particle size distribution of AuNPs obtained from the TEM image analysis. The data were fitted to the log-normal distribution. The peak values of the fit were taken as the average particle size of the corresponding samples. D. UV-Vis-Absorption Spectra of AuNP and AuNS demonstrate absorbance peaks. E. Zeta-potential measurement.

### Various degrees of toxicity of AuNP and AuNS to *C. reinhardtii*

To evaluate the impact of different shapes of Si-AuNPs on the growth of *C. reinhardtii*, and to determine whether there are differences in toxicity between various concentrations of AuNP and AuNS, we measured the growth rate by recording optical density (OD) values every 24 h for 5 days. The results demonstrated a significant reduction in growth rate compared to the control group following exposure to any shape and concentration of AuNPs for 48-120 h, with more severe inhibition of *C. reinhardtii* growth observed at higher concentrations of AuNPs. Notably, at the same concentration, AuNS caused greater growth inhibition than AuNP at each time point. Statistically significant differences were observed between the 10 mg/L AuNP and AuNS groups at 72, 96, and 120 h, and between the 1 mg/L AuNP and AuNS groups at 120 h (Figure 2A).

**Figure 2.**
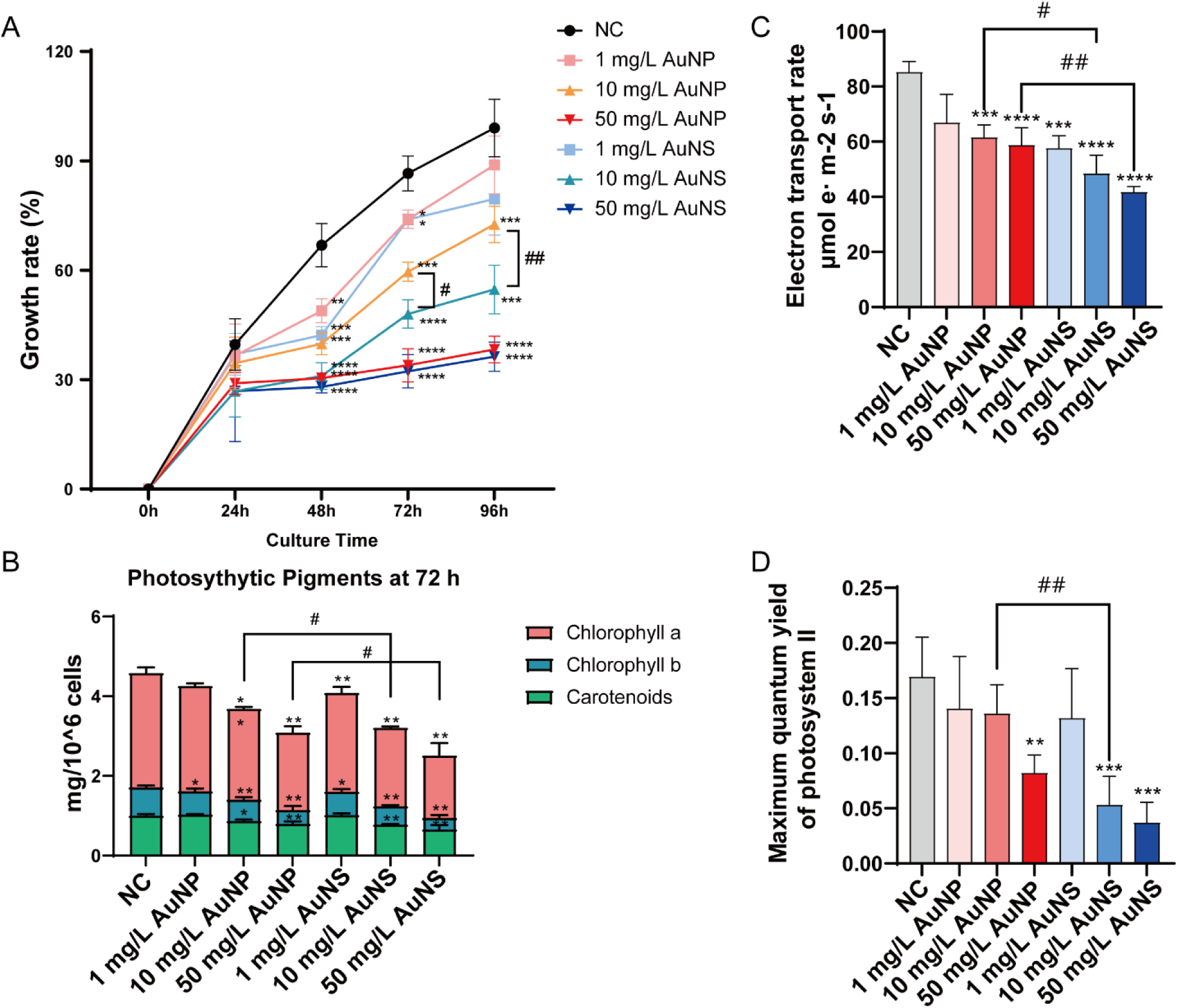
Different hazard effect of AuNP and AuNS on *C. reinhardtii*. A. Growth rate from 24 to 120 h. B. Concentration of photosynthetic pigments at 72 h. C. Electron transport rate at 72 h. D. Maximum quantum yield of photosystem II at 72 h. Data are presented as mean ± SD. n = 3. * *p* < 0.05, ** *p* < 0.01, *** *p* < 0.005, and **** *p* < 0.001 compared with control group or between the two groups with lines.

To further verify the impact of AuNPs with different shapes on *C. reinhardtii*, we measured the concentrations of three main photosynthetic pigments, namely chlorophyll a, chlorophyll b, and carotenoids. Due to the significant differences in growth inhibition observed between the 10 mg/L AuNP and AuNS groups at 72 h, we selected 72 h as the effective time point for measuring photosynthetic pigments concentrations. As results shown in Figure 2B, chlorophyll a exhibits significant decreases across all exposure groups compared to the control group, except for the 1 mg/L AuNP group. Chlorophyll b shows significant reductions across all concentrations, while carotenoids display a significant decline in the 10 and 50 mg/L groups for both AuNP and AuNS. These results indicate that higher concentrations of AuNPs lead to more pronounced inhibitory effects on the photosynthetic pigments of *C. reinhardtii*. Similar to the growth rate findings, exposure to AuNS results in a more pronounced inhibitory effect on pigments compared to AuNP, with statistically significant differences observed in the chlorophyll a concentration in both the 10 and 50 mg/L dosage groups.

We also assessed the ETR and Y(II), across all AuNPs concentrations groups at 72 h. Both parameters were reduced compared to the control group (Figure 2C and 2D), indicating the inhibitory effect of AuNPs on algal photosynthetic efficiency. Statistically significant differences in ETR were observed between the 10 and 50 mg/L AuNP and AuNS groups, while Y(II) showed differences between the 10 mg/L groups. Therefore, exposure to 10 mg/L AuNPs for 72 h appears to be an effective concentration and a sensitive time point for assessing toxicity.

### Effects of AuNP and AuNS exposure on *C. reinhardtii*: size, granularity, and debris production

Figure S1A shows FSC and SSC signals of algae cells following exposure to AuNPs with different shapes. In the control group, microalgae cells appear as homogeneous and elliptical populations centered in the plot (56.24% in the P3 region), indicating no cell aggregation, morphological changes, or damage. Debris was observed in the bottom left corner. Upon exposure to AuNPs, the proportion of events in the P3 region decreased to 51.25%, 50.37%, and 46.39% for the 1, 10, and 50 mg/L AuNP groups, and to 47.91%, 37.93%, and 32.51% for the 1, 10, and 50 mg/L AuNS groups, respectively. Significant differences were observed in the 50 mg/L AuNP group and the 10 and 50 mg/L AuNS groups compared to the control (Figure S1B). Interestingly, significant changes in the shape and position of the main populations (P3) were noted in the 10 and 50 mg/L AuNS groups, which we designated as the P2 region. Events in P2 accounted for 48.19% and 40.49% of the total events, respectively. Additionally, after exposure to AuNPs, there was an emergence of events in the left side (P4), interpreted as debris resulting from cell damage and death, with numbers increasing with AuNPs concentration. Similar to P3, there were significant differences in the 50 mg/L AuNP group and the 10 and 50 mg/L AuNS groups compared to the control. Furthermore, the percentage of debris was significantly different between the 50 mg/L AuNP and AuNS groups (Figure S1C). To rule out the influence of nanoparticles themselves on these results, we conducted FC experiments with different concentrations of AuNPs solutions only (Figure S2), which confirmed that debris in P4 was not affected by AuNPs concentration. Thus, exposure to AuNPs result in increased cell size, enhanced internal complexity, and greater amounts of cellular debris following cell damage or death. Notably, differences in the proportion, shape, and localization of P3/P2 and P4 events between AuNP and AuNS groups indicate varying impacts of differently shaped AuNPs on microalgae, with those exposed to AuNS displaying larger cell bodies, more complex internal granularity, and increased debris.

Given that the FC results indicated cell enlargement and increased debris upon exposure to AuNPs, we conducted further observations of cell morphology and quantified the major axis and cell area using TEM. We selected 10 and 50 mg/L AuNPs exposure groups at 72 h, as the 1 mg/L groups showed negligible effects on the microalgae. As depicted in Figure S3A, S3B, and S3C, both the major axis and the area of the cells in the AuNPs exposure groups were significantly larger than those in the control group, with an increase observed as the concentration rose. Notably, at equivalent exposure concentrations, cells in the AuNS groups were larger than those in the AuNP groups. This difference was particularly significant for cell area in the 50 mg/L group. To further investigate the increased granularity observed, we examined the subcellular structure of *C. reinhardtii* to confirm the internal structural changes.

### Different accumulation and disruption of algal subcellular structures by AuNPs

Firstly, we employed confocal laser scanning microscopy (CLSM) to examine the accumulation of AuNPs in *C. reinhardtii* after 72 hours of exposure (Figure 3A). AuNPs were labeled with Alexa Fluor 488 (AF488) dye to determine their localization and accumulation within the algal cells. Chlorophyll autofluorescence (magenta) indicated the position of chloroplasts, while AuNPs were visualized via green fluorescence from the AF488 label. Manders’ colocalization coefficient analysis was performed on images displaying fluorescent signals from AF488-labeled AuNPs and chloroplast autofluorescence.

**Figure 3.**
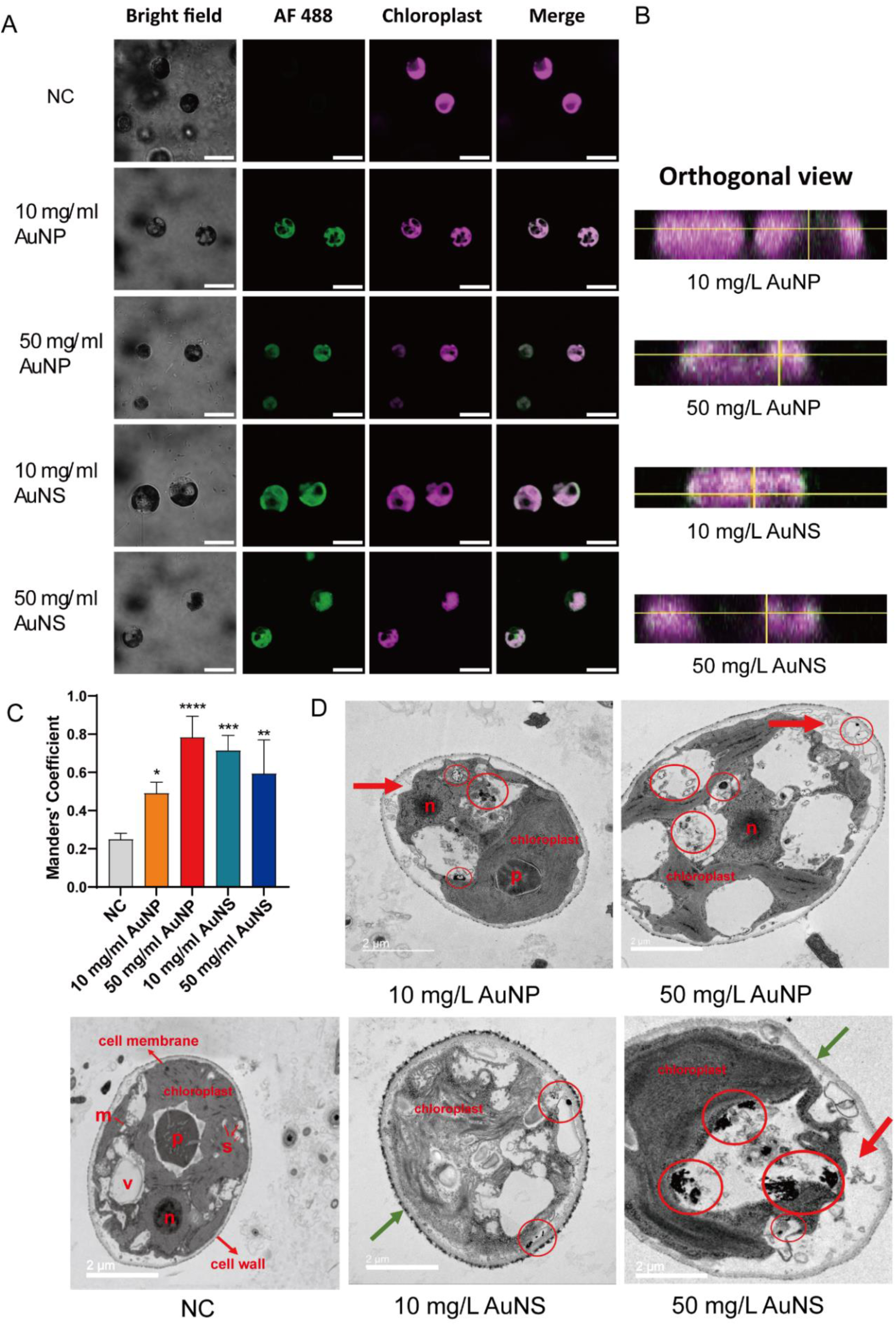
Interaction between *C. reinhardtii* and AuNPs after 72 h exposure. A. Representative confocal images of chloroplast and AuNPs (Scale bar: 10 μm). B. Orthogonal view of confocal images. C. The degree of colocalization between two channels in confocal images. D. Representative TEM images of microalgae and AuNPs interaction. Scale bars: 2 μm. Subcellular structures are: n, nucleus; p, pyrenoid; v, vacuole; m, mitochondrion; s, starch granule. Red circles indicate AuNPs entering the cell and aggregating in vacuole and cytoplasm, red arrows indicate the plasmolysis, and green arrows indicate cell wall thicken.

Both types of nanoparticles, spherical (AuNP) and star-shaped (AuNS), were internalized by the algal cells, with their localization confirmed in the chloroplasts. Overlay images of AF488 (green) and chloroplasts (magenta) revealed white regions in the merged channels, indicating colocalization at both 10 mg/L and 50 mg/L concentrations for AuNP and AuNS. Orthogonal projections of confocal images (Figure 3B) showed that a portion of the AuNP uniformly adhered to the surface of the chloroplast membrane, forming a thin green fluorescent ring surrounding the magenta chloroplast autofluorescence.

Manders’ colocalization coefficients between AuNP (AF488) and chloroplasts (Figure 3C) were approximately 0.5 at 10 mg/L and 0.76 at 50 mg/L, both significantly higher than the negative control (NC) group. In contrast, the green fluorescence of AuNS was not evenly distributed within the chloroplasts. Instead, AuNS tended to aggregate on the chloroplast surface, forming bright fluorescent spots, or they penetrated the chloroplast membrane and accumulated in other intracellular compartments (Figure 3A), a pattern also evident in the orthogonal projections (Figure 3B). Notably, statistical analysis (Figure 3C) showed that in the 10 mg/L AuNS group, the colocalization coefficient was ∼0.73, similar to that of the 50 mg/L AuNP group. However, in the 50 mg/L AuNS group, the colocalization coefficient decreased to ∼0.6. This reduction may be attributed to the sharp angular features of AuNS, which likely facilitate membrane puncture, leading to partial escape from the chloroplasts and subsequent accumulation in other intracellular regions.

Then we used TEM to scan the subcellular structures of *C. reinhardtii*. As Figure 3D showed that, in the control group, the morphology of *C. reinhardtii* cells was intact, with the cell membrane closely adhering to the cell wall and no signs of plasmolysis. The internal structures, including the chloroplasts, pyrenoids, mitochondria, and starch granules, were intact, and the vacuoles were clear and free of impurities. Upon exposure to 10 mg/L of AuNP, the cell structure remained largely intact, but signs of plasmolysis were evident. AuNP were observed within the cytoplasm and vacuoles. In the 10 mg/L AuNS group, the cell structure showed mild disruption, with the absence of pyrenoids, fewer starch granules in the chloroplasts, and a thickened cell wall. AuNS were observed in the wall membrane space, cytoplasm, vacuoles, and other organelles. Further, cells exposed to 50 mg/L of AuNP resulted in severe structural damage. Significant plasmolysis was observed, accompanied by the absence of pyrenoids and a reduction in starch granules. Numerous AuNP were found within the cells. In the 50 mg/L AuNS exposure group, cellular aberrations were evident. The normal cell structure was almost entirely disrupted, with severe plasmolysis, no pyrenoids or starch granules, a thick and blurred cell wall, and a substantial accumulation of AuNS throughout the cells.

### Effect of AuNP and AuNS on algal gene expression profiles and function enrichment

Given the distinct types of damage observed in *C. reinhardtii* after exposure to AuNP and AuNS, we hypothesize that the varying shapes of AuNPs leads to differential gene expression and distinct injury mechanisms. Considering the severe cellular damage noted in the 50 mg/L AuNPs groups and the significant toxic effects observed in the 10 mg/L exposure groups between AuNP and AuNS, we conducted RNA-seq to analyze gene expression in microalgae exposed to 10 mg/L AuNP and AuNS for 72 h. The raw data required to reproduce the above findings are available to download from (Reserved DOI: 10.17632/x3gyskppmp.1). Quality control (QC) for raw data is presented in Table S1. A total of 15,087 genes were identified when comparing the NC vs. AuNP groups, with 9 genes upregulated and 38 downregulated. In contrast, the NC vs. AuNS comparison revealed 246 upregulated and 145 downregulated genes among the 15,175 detected genes (Figure 4A). There were 34 differentially expressed genes (DEGs) co-differentially expressed in both the AuNP and AuNS exposure groups (Figure 4B). The top 18 major Gene Ontology (GO) terms (*p* < 0.05) are presented, including ten terms related to biological processes and eight terms related to cellular components identified under 10 mg/L AuNP and AuNS exposure groups.

**Figure 4.**
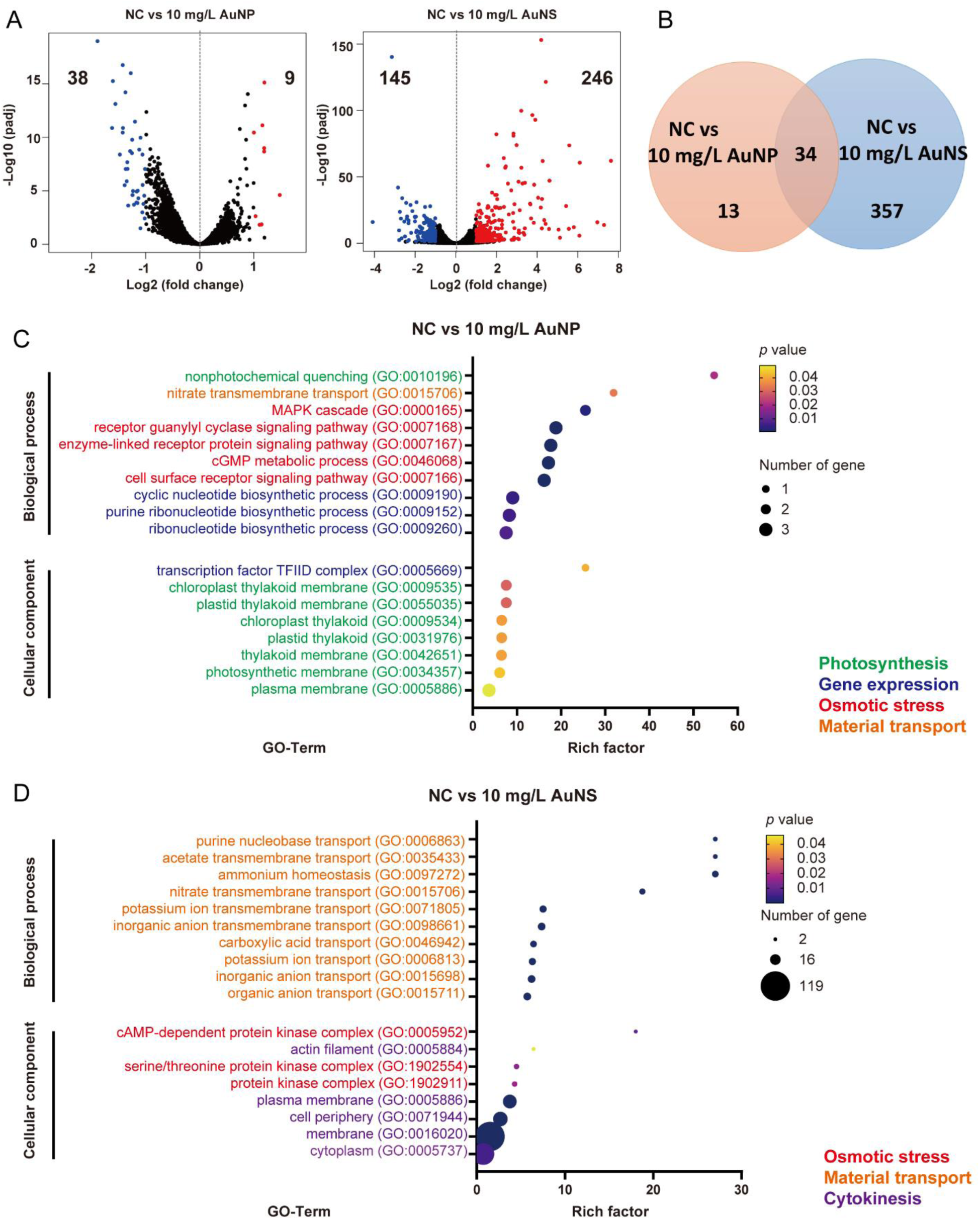
Exposure to AuNP and AuNS caused *C. reinhardtii* gene expression and functional enrichment changed. A. Volcano plots of quantified genes in the 10mg/L AuNPs exposed group as compared to the control group. Significant increased genes are highlighted in red and decreased genes are in blue (*p* < 0.05, fold change > 1 or < -1). B. Venn diagram showing the number of the same DEGs and unique DEGs between the AuNP and AuNS exposure groups. Gene Ontology (GO) analysis of DEGs pathways in (C) 10 mg/L AuNP and (D) AuNS treatment, respectively. The up 10 pathways are biological process and down 8 pathways are molecular function, ranked by number of rich factors. Terms related to photosynthesis are in green, gene expression is in blue, osmotic stress are in red, material transport are in orange, cytokinesis are in purple.

In the NC vs AuNP group, DEGs were associated with various aspects of photosynthesis, gene expression regulation, osmotic stress processes, and material transport. The most enriched term with the highest rich factor among biological processes was non-photochemical quenching (NPQ), which contributes to photosynthesis. Seven of the eight cellular component terms were involved in photosynthesis, including the chloroplast thylakoid membrane, plastid thylakoid membrane, chloroplast thylakoid, plastid thylakoid, thylakoid membrane, photosynthetic membrane, and plasma membrane. Biological processes, including the mitogen-activated protein kinase (MAPK) cascade, receptor guanylyl cyclase signaling pathway, enzyme-linked receptor protein signaling pathway, guanosine 3’,5’-cyclic monophosphate (cGMP) metabolic process, and cell surface receptor signaling pathway, were implicated in the regulation of osmotic stress. Gene expression regulation processes, such as cyclic nucleotide biosynthetic process, purine ribonucleotide biosynthetic process, and ribonucleotide biosynthetic process, along with the cellular component transcription factor II D (TFIID) complex, were identified. Nitrate transmembrane transport was associated with material transport (Figure 4C).

In the NC vs AuNS group, DEGs participated in different functions. In addition to osmotic stress processes (cellular component: cAMP-dependent protein kinase complex, serine/threonine protein kinase complex, and protein kinase complex), all biological process terms related to material and transmembrane t/ransport. This included purine nucleobase transport, acetate transmembrane transport, ammonium homeostasis, nitrate transmembrane transport, potassium ion transmembrane transport, inorganic anion transmembrane transport, carboxylic acid transport, potassium ion transport, inorganic anion transport, and organic anion transport. As well as five cellular component terms related to cytokinesis, including actin filament, plasma membrane, cell periphery, membrane, and cytoplasm (Figure 4D).

### Differential toxic mechanisms of AuNP and AuNS effects on *C. reinhardtii*

Figure 5A illustrates the expression profiles of the 34 co-differentially expressed genes (co-DEGs) across the control, AuNP, and AuNS groups. The patterns of upregulation or downregulation were consistent within the two groups; however, the fold changes were more pronounced in the AuNS group compared to the AuNP group. This observation aligns with the results on cell growth inhibition, morphological alterations, and subcellular structural damage, indicating that the cytotoxicity induced by AuNS was consistently more severe than that caused by AuNP. Despite this, STRING analysis of these 34 genes revealed no functional connections (Figure S4), suggesting that the co-DEGs might not contribute significantly to the primary mechanism of toxicity.

**Figure 5.**
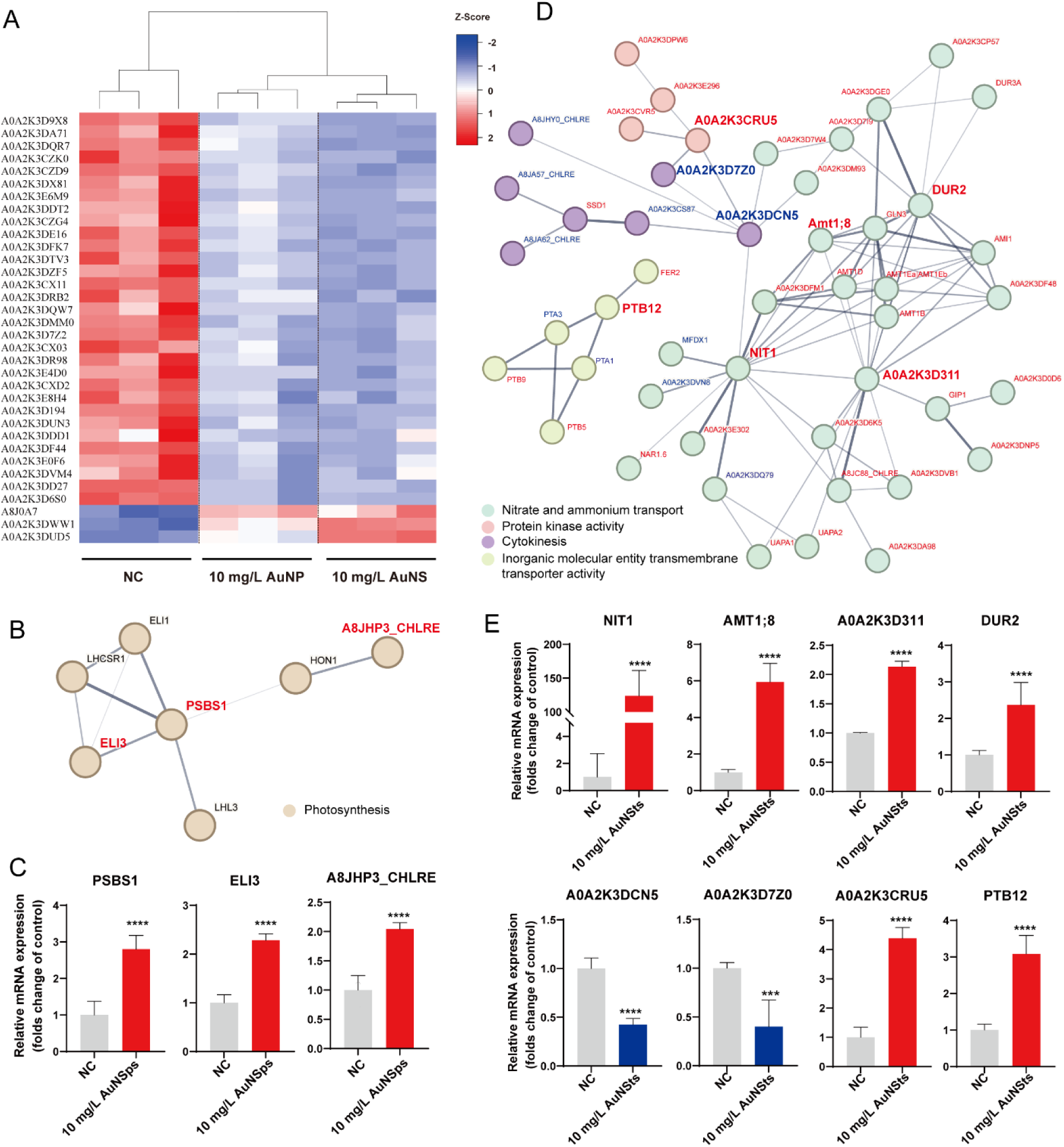
Exposure to AuNP and AuNS caused different toxic mechanisms through the individually differential expression genes. A. Heat map of co-DEGs in the control group, 10 mg/L AuNP and 10 mg/L AuNS exposure groups. Significant increased genes are highlighted in red and decreased genes are in blue (*p* < 0.05, fold change > 1 or < -1). B. String analysis of individually differential expression genes in 10 mg/L AuNP group. The edges indicate both functional and physical protein associations. Line thickness indicates the strength of data support. C. mRNA expression levels of key genes in 10 mg/L AuNP group (n = 3). D. String analysis of individual differential expression genes in 10 mg/L AuNS group. E. mRNA expression levels of key genes in 10 mg/L AuNS group (n = 3). The genes in the STRING analysis graph that were marked with the same color represent potential interactions or regulatory relationships among them. Edges represent protein-protein associations are meant to be specific and meaningful, i.e. proteins jointly contribute to a shared function; this does not necessarily mean they are physically binding to each other. Data are presented as mean ± SD. *** *p* < 0.005 and **** *p* < 0.001 compared with control group.

Subsequently, we conducted functional connectivity analysis using STRING for genes individually altered in either the AuNP or AuNS groups. In the AuNP group, STRING analysis of 13 individually altered genes, along with four introduced interactors, revealed interactions among three genes forming a cluster centered around the photosystem II subunit S1 (PSBS1), with early light-inducible protein (ELI3) and A8JHP3 also showing increased expression (Figure 5B and 5C). In the AuNS group, STRING analysis of 357 individually altered genes, excluding non-interacting ones, identified four prominent clusters (Figure 5D). Key genes within these clusters, identified based on their interaction strength and functional relevance, included nitrate reductase (NIT1), aminomethyl transferase (AMT1), uricase (A0A2K3D311), urea amidolyase (DUR2), RING-type domain-containing protein (A0A2K3DCN5), Cyclin-like domain-containing protein (A0A2K3D7Z0), protein kinase domain-containing protein (A0A2K3CRU5), and phosphate transporter (PTB12).

## Discussion

The inhibition of algal growth can be primarily attributed to two factors: (1) NPs disrupt the photosynthetic process in microalgae by blocking or scattering light depending on their sizes, thereby reducing the efficiency of photosynthesis and impacting the microalgae’s energy production capabilities^45^; (2) chemical damage: NPs damage cell membranes by inducing excessive accumulation of reactive oxygen species (ROS), which in turn impairs chlorophyll synthesis, thereby diminishing the microalgae’s capacity to absorb and convert light energy^46^. The initial impact of these damages is the disruption of photosynthesis, given that photosynthetic pigments are crucial for capturing sunlight energy for photosynthesis in algae^47^. Accordingly, our results indicate that the growth rate of *C. reinhardtii* decreased upon exposure to both AuNP and AuNS across all concentrations, with photosynthetic pigment inhibition. Monitoring of photosynthesis is a good approach to gain general information on plant performance in growth and reproduction^48^. Both ETR and Y(II) decreased compared to the control group, signifying an inhibitory effect of AuNPs on algal photosynthetic efficiency existed. The reduced values of ETR and Y(II) indicate plant stress as they are indicators of photosynthetic activity^43^. Notably, all these results show statistically significant differences between the 10 mg/L AuNP and AuNS groups, with AuNS exhibiting a more severe inhibitory effect. It can be inferred that star-shaped AuNPs were more damaging to microalgae than their spherical counterparts.

Our flow cytometry results show that post-exposure to AuNPs, *C. reinhardtii* cells exhibit increased size, enhanced internal complexity, and a greater quantity of cellular debris, indicative of cellular damage or death. Notably, the events proportion, shape, and localization of P4 area in Figure 3A varied between AuNP and AuNS groups, suggesting that the shape of AuNPs distinctly impacts microalgae. Specifically, cells exposed to AuNS display larger cell bodies, more complex internal granularity, and increased debris compared to those exposed to AuNP. TEM images at low magnification further confirmed that exposure to AuNS caused larger cell volumes than exposure to AuNP. It is well-established that NPs can inhibit photosynthesis in microalgae by blocking or scattering light, leading to reduced photosynthetic efficiency, cellular swelling, and increased debris production^45^. Thompson et al. found that *C. reinhardtii* strains dramatically swell and ultimately disintegrate when grown in the dark, which can result in the leakage of cellular contents due to the destruction of the cell wall and membrane from swelling^49^. This phenomenon may explain the observed cell enlargement, increased internal complexity, and elevated debris levels.

NPs can physically interact with microalgae cells and be taken up from the surrounding medium, this process may block or damage cell membranes, thereby compromising cellular integrity and function^45^. Plasmolysis observed in *C. reinhardtii* post AuNPs exposure might be attributed to this mechanism. AuNS may more easily traverse cellular barriers, potentially exacerbating damage due to their structural characteristics^50^. Once internalized, NPs can accumulate within various organelles and subcellular structures, disrupting normal cellular functions^45^. Most cells exposed to AuNPs exhibited reduced internal structures, with significant membrane deformations and absence of clearly identifiable vacuoles compared to control groups. Specifically, cellular structures in the AuNS groups were notably more disorganized than those in the AuNP groups at equivalent exposure concentrations. Moreover, the severity of internal structural damage was most pronounced in the 50 mg/L groups. The occurrence of chloroplasts containing starch granules and the central pyrenoid reveals high photosynthetic activity and energy stock capacity ^51^. Notably, AuNS (10 mg/L) and AuNP (50 mg/L) resulted in fewer starch granules and a reduced occurrence of pyrenoid, indicating that lower AuNS concentration might affect photosynthetic and metabolic activity in *C. reinhardtii*. Cell wall thickening is one of the primary indicators of nitrogen deficiency^52^. Nitrogen deprivation can result from nutrient deprivation or changes in light intensity^53^. This may explain the observed cell wall thickening in the AuNS group. In addition, nitrogen limitation affects the growth of microalgae^54^, which also accounts for the lower growth rate of cells in the AuNS group compared to the AuNP group. Through cytotoxicity assays and observations of cell morphology and subcellular structures, we conclude that AuNS exhibit greater toxicity towards *C. reinhardtii* compared to AuNP.

The 10 mg/L concentration allowed us to detect clear differences in internalization behavior and subcellular injury between AuNP and AuNS. So that we selected this concentration for further mechanism investigation. Based on RNA-seq analysis, DEGs in NC vs AuNP group and NC vs AuNS group enriched in distinct biological processes and cellular components. It has been shown that AuNPs in leaves can lead to fluorescence quenching, as the excited electron of chlorophyll molecules can be transferred to AuNPs, inducing the increase of NPQ in plants^55^. Saison et al. found that copper oxide NPs covered with polystyrene had a deteriorative effect on chlorophyll by inducing the photoinhibition of photosystem II, and the inhibition of photosynthetic electron transport induced a strong dissipation of energy by NPQ processes^56^. Furthermore, the reorganization of the photosynthetic machinery, especially in the range of alternative energy-dissipation pathways has been reported as responses to osmotic stress^57^. Our GO enrichment analysis indicate that AuNP primarily affected photosynthesis and osmotic stress in *C. reinhardtii*, resulting in inhibited cell growth, cell death, and damage to subcellular structures. In contrast, AuNS primarily affected material transport and cytokinesis as their main toxic mechanisms. Substance transport and transmembrane transport are involved in all biological processes, primarily concerning nitrogen and related metabolites, such as purine^58^ and ammonium^59^, which is the main reason for growth inhibition of microalgae. The actin filaments, plasma membrane, cell periphery, membrane, and cytoplasm play crucial roles in cytokinesis^60^. Therefore, despite AuNS appearing more severe in their effects on microalgae compared to AuNP, the underlying mechanisms differ significantly.

According to our co-DEGs and individually DEGs results, the toxic effects of AuNP on *C. reinhardtii* appear to stem from alterations in specific genes rather than co-DEGs. In the AuNP group, the cluster’s main biological process, molecular function, and cellular component were non-photochemical quenching, chlorophyll binding, and photosystem II, respectively. These findings align closely with the GO analysis results for the AuNP group (Figure 4C), suggesting that the toxic effects of AuNP on *C. reinhardtii* may primarily result from alterations in these individually affected genes rather than the co-DEGs. PSBS1 plays a pivotal role in NPQ, where its expression increases during NPQ to modulate chlorophyll fluorescence quenching in photosystem-depleted membranes^61^. ELI3, located in thylakoid membranes, is known to protect photosynthetic machinery from various environmental stresses in plants^62^, with its expression heightened under photo-oxidative stress conditions^63^. A8JHP3 is the major protein complex of the chloroplast photosynthetic apparatus, and the increased expression also contributes to abiotic stress tolerance^64^. The elevated expression of these three genes in the AuNP exposure group suggests that PSBS1/ELI3 pathway is pivotal in impairing the photosynthetic capacity of *C. reinhardtii*, thus serving as the primary toxic mechanism induced by AuNP exposure.

In the AuNS group, the largest cluster was associated with nitrate and ammonium transport, encompassing processes such as ammonium homeostasis, nitrate assimilation, and ammonium transmembrane transporter activity. The other three clusters pertained to protein kinase activity, cytokinesis (involving plasma membrane and cell periphery), and inorganic molecular entity transmembrane transporter activity (including cellular responses to inorganic substances). These functions of the individually altered genes in the AuNS group are consistent with the GO analysis results (Figure 4C). NIT1 functions as a crucial enzyme involved in the initial step of nitrate assimilation in plants^65^. Nitrate reductases, classified as molybdoenzymes, catalyze the reduction of nitrate to nitrite, a pivotal reaction in protein production in most crop plants, where nitrate serves as the primary nitrogen source in fertilized soils. AMT play a central role in the uptake of reduced nitrogen for biosynthetic and energy metabolism^66^. A0A2K3D311 is essential in ureide formation^67^, while DUR2 catalyzes urea conversion to ammonium, the key step in utilizing urea as a nitrogen source^68^. The elevated expression of these four genes represented increased nitrogen metabolism and transport.

Although previous studies have suggested that AuNPs could contribute to the enhanced nitrogen metabolism in plants^69^, our experimental results indicate that the upregulation of nitrogen metabolism-related genes observed here is likely compensatory. This compensatory regulation may result from significant energy loss caused by photosynthetic inhibition, prompting microalgae to rely more on nitrogen metabolism to bolster nutrition. Functional studies of A0A2K3DCN5 and A0A2K3D7Z0 are currently limited; however, Uniport UniProt annotations indicate that the gene cluster in which they serve as key components are associated with cytokinesis. Thus, their reduced expression may reflect impaired cytokinesis and, consequently, decreased cell proliferation. A0A2K3CRU5 is a structurally conserved protein domain containing the catalytic function of protein kinases^70^. These enzymes can move a phosphate group onto proteins, inducing phosphorylation^71^. Protein phosphorylation is a reversible modification that plays a crucial role in plant signaling transduction and environmental adaptation^72^. The exposure to the foreign substances, such as AuNS, may have stimulated a cellular response leading to the upregulation of genes within this cluster. PTB12 plays a fundamental housekeeping role in phosphate transport^73^, critical for plant phosphate homeostasis mediated by membrane-localized transporters^74^. Dysregulated expression of genes in this cluster suggests membrane transport disturbances possibly induced by AuNS compromising membrane integrity. The aberrant expression patterns of these genes in the AuNS exposure group imply multiple pathways of damage, particularly affecting nutrient metabolism, transmembrane transport, and cell proliferation. The NIT1/AMT1/A0A2K3CRU5 pathway emerges as a potential primary toxic mechanism induced by AuNS exposure.

Our findings indicate unique mechanisms by which different shapes of AuNPs interact with *C. reinhardtii*. Specifically, AuNP predominantly impact photosynthesis and induce cellular damage, whereas AuNS primarily disrupt cell membrane structure, impairing transmembrane transport and substance absorption. These effects can be attributed to their respective shapes: AuNP, being spherical with smooth surfaces, are prone to adhere to the surfaces of microalgae cells and chloroplast membranes. The adherence of AuNP of this size likely causes light scattering, attenuating photosynthesis, leading to NPQ, cell swelling, nutrient deprivation, and eventually cell death. In contrast, AuNS, characterized by their star-shaped and irregular protrusions, are more likely to penetrate cell membrane structures. This penetration results in extensive membrane damage, hindering transmembrane transport of materials, causing cell rupture, disintegration of internal structures, and cell death. However, the precise interactions between different shapes of AuNPs and cells require further investigation. Future studies could benefit from employing artificial intelligence to simulate the motion and interaction pathways of these nanoparticles, offering deeper insights into their cellular dynamics.

## Conclusion

our study demonstrates that AuNS exert more severe effects on *C. reinhardtii* cells compared to AuNP, including growth inhibition, photosynthesis suppression, cell enlargement, debris production, and subcellular structure disruption. This heightened toxicity can be attributed to the distinct mechanistic interactions driven by the nanosphere and nanostar morphologies of the nanoparticles. AuNP tend to adhere to the membrane surfaces of *C. reinhardtii* through occlusion, leading to diminished cellular photosynthesis and reduced energy storage via the PSBS1/ELI3 pathway, which results in moderate cell damage and death. In contrast, AuNS compromise membrane structure and disrupt cellular metabolism through the NIT1/AMT1/A0A2K3CRU5 pathway, severely affecting the physiological functions of *C. reinhardtii* and causing extensive cell damage and death. This study elucidates the unique toxic mechanisms by which different nanoparticle shapes interact with microalgae, underscoring the importance of nanoparticle shape in assessing ecotoxicity. Our findings suggest that in the development and production of nanoparticles, consideration should extend beyond application performance stability and functional enhancements to include potential ecological and human health impacts. Emphasizing nanoparticle morphology optimization to maintain safety and stability while preserving functionality is essential for minimizing adverse environmental effects.

## Acknowledgements

This work was supported by NSF CBET (Project No. 2035623 and No. 2034855) and NIH (Grant No. R35GM 142763). We express our gratitude to UCSD Cellular and Molecular Medicine Electron Microscopy Core (UCSD-CMM-EM Core, RRID: SCR_022039) for their assistant on TEM imaging. Dr. Can Wang thanks to Fatemeh “Sara” Nafar for her assistance in training on microalgae cell culture techniques.

## Notes

The authors declare no competing financial interest.

## Supporting Information

RNA-seq based exposure concentration raw data, experimental results, and additional results are presented in the supplemental materials.

## Author Contributions

**Can Wang**: Conceptualization, Investigation, Methodology, Visualization, Writing original draft. **Bhaskar Sharma:** Methodology, Data Analysis, Writing – review. **Ruonan Peng**: Methodology, Data Analysis. **Xin Yong**: Supervision, Validation. **Louis S. Santiago**: Methodology. **Ke Du**: Conceptualization, Supervision, Writing - review & editing.

---

**Table S1.**
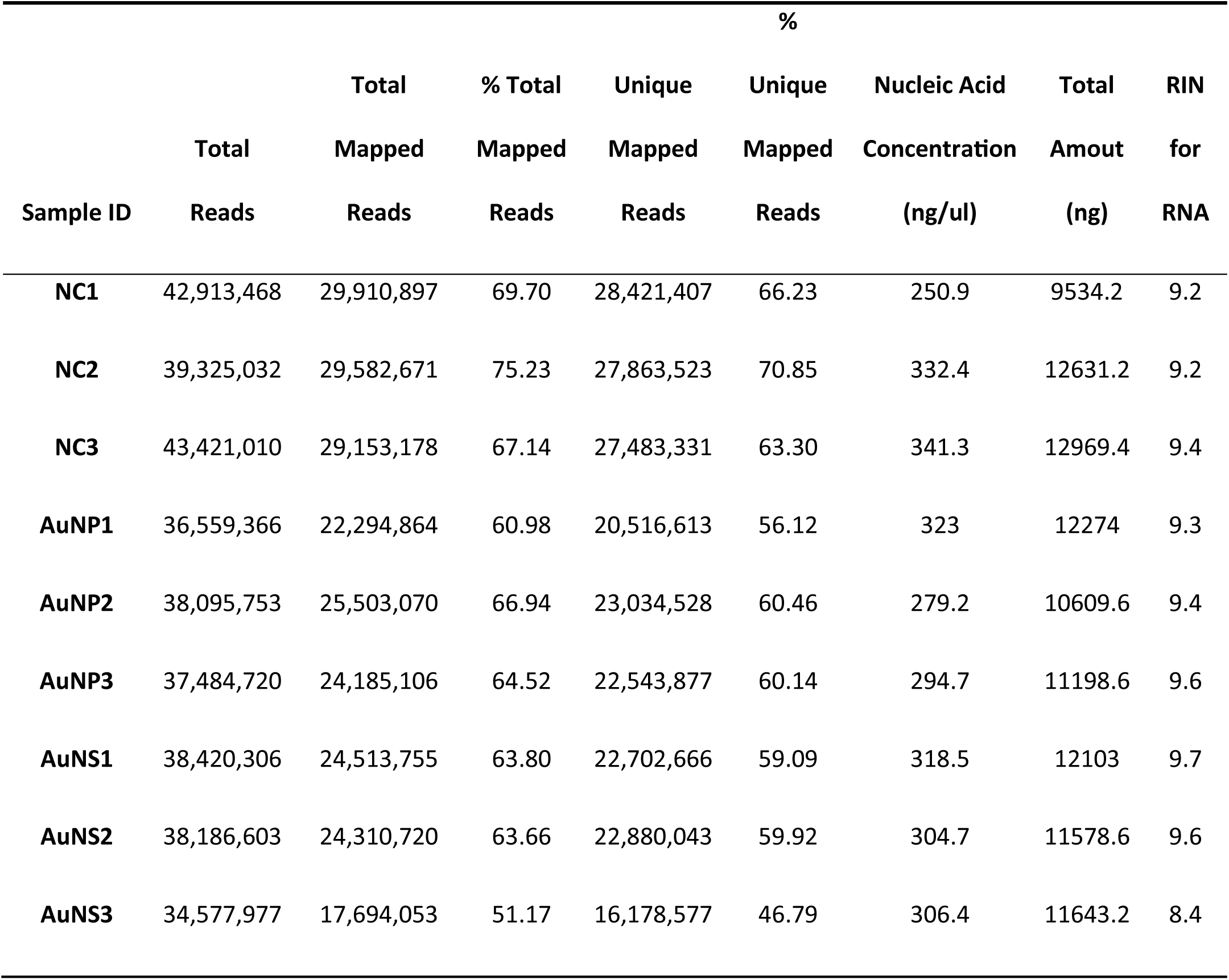
RNA-Seq Quality Control (n=3)

**Figure S1.**
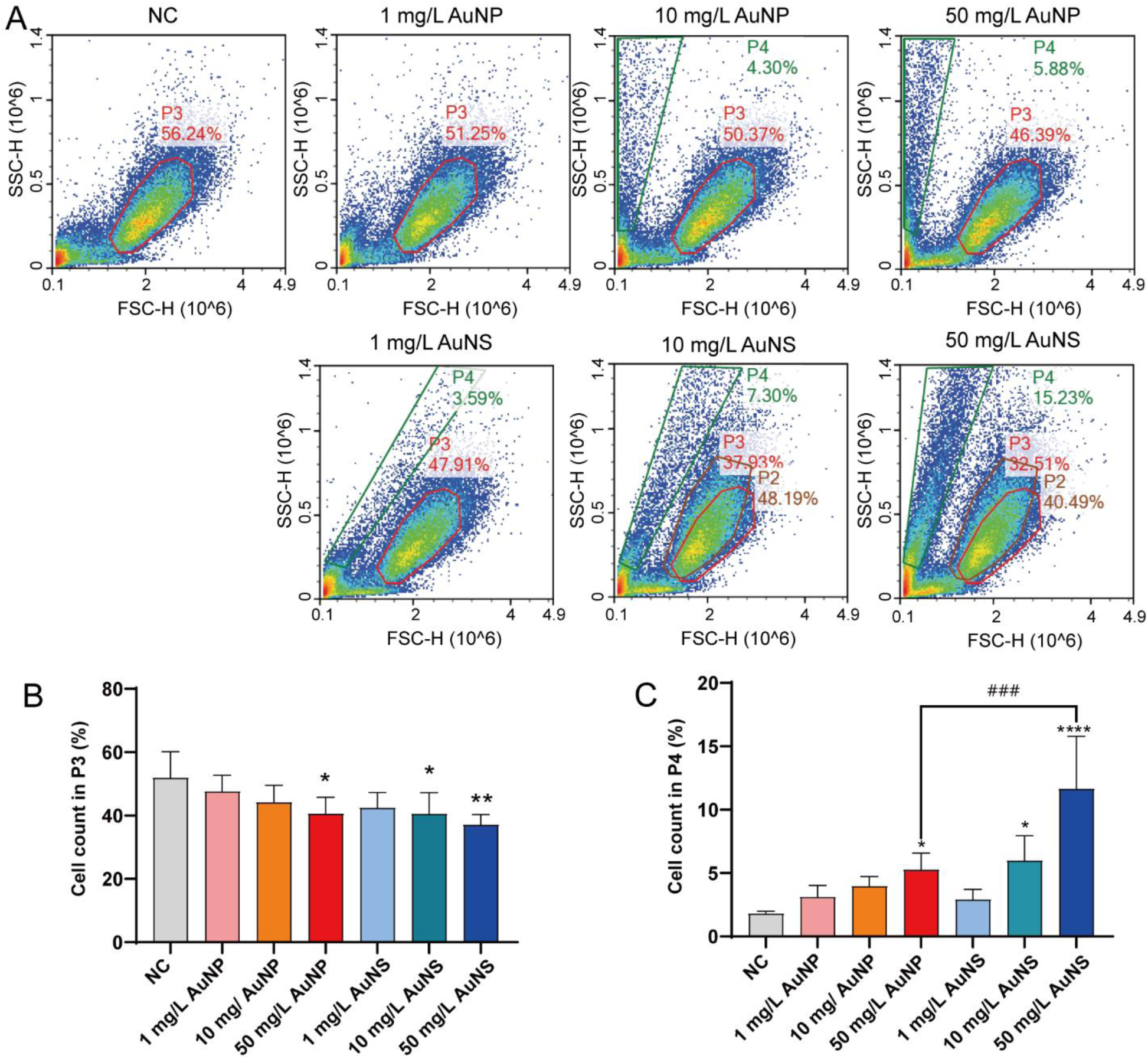
Exposure to AuNP and AuNS caused *C. reinhardtii* producing debris with different shape and granularity as well as different cells size and shape. A. The flow cytogram of forward scatter height (FSC-H) vs. side scatter height (SSC-H). B. Number of live cells located within the P3 gate. C. Amount of debris within the P4 gate. Data are presented as mean ± SD. n = 3. * *p* < 0.05, ** *p* < 0.01, *** *p* < 0.005, and **** *p* < 0.001 compared with control group or between the two groups with lines.

**Figure S2.**
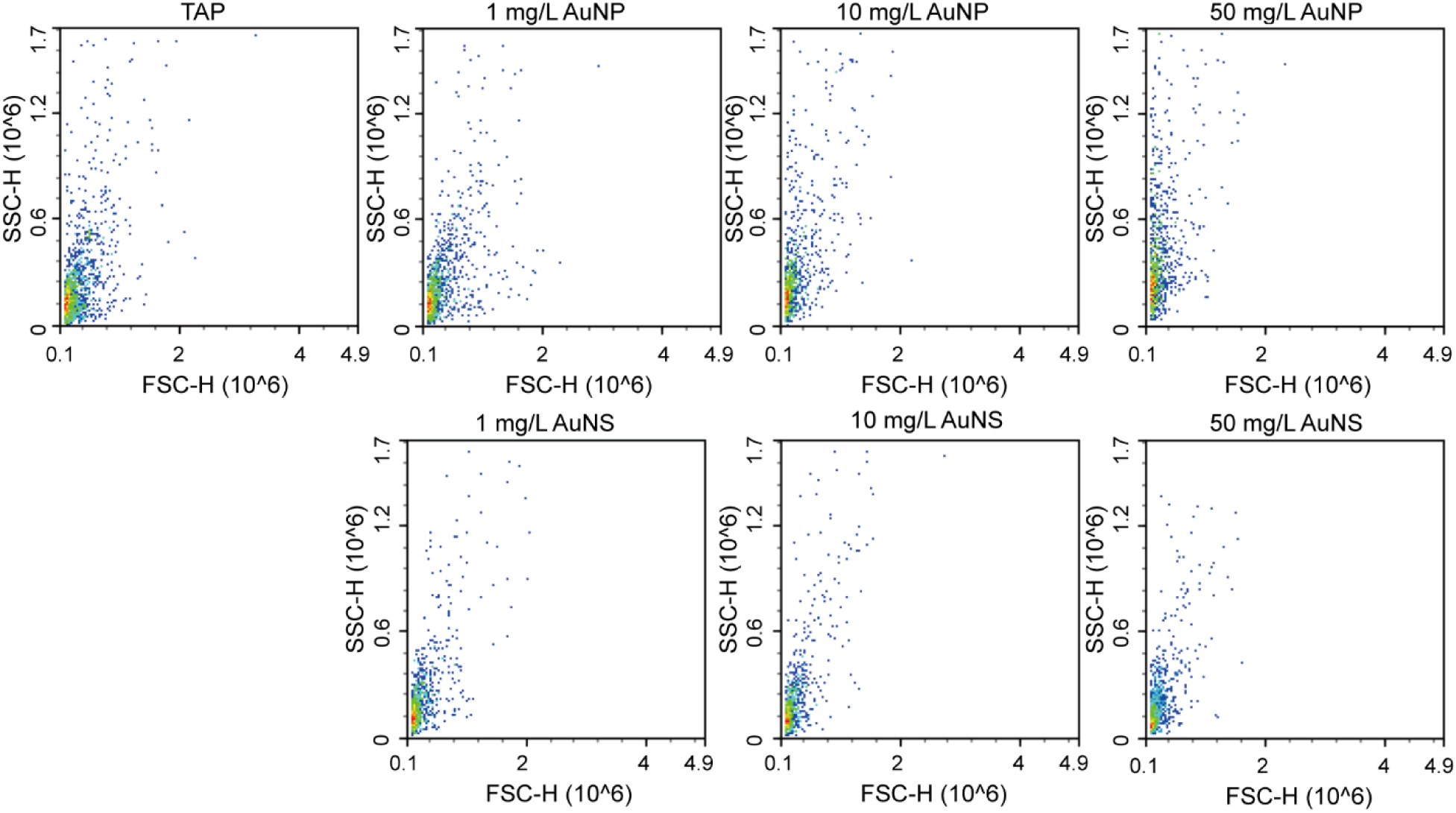
FC experiments with different concentrations of AuNPs solutions. The flow cytogram of forward scatter height (FSC-H) vs. side scatter height (SSC-H) of cells in NC and AuNPs treatment groups (n=3).

**Figure S3.**
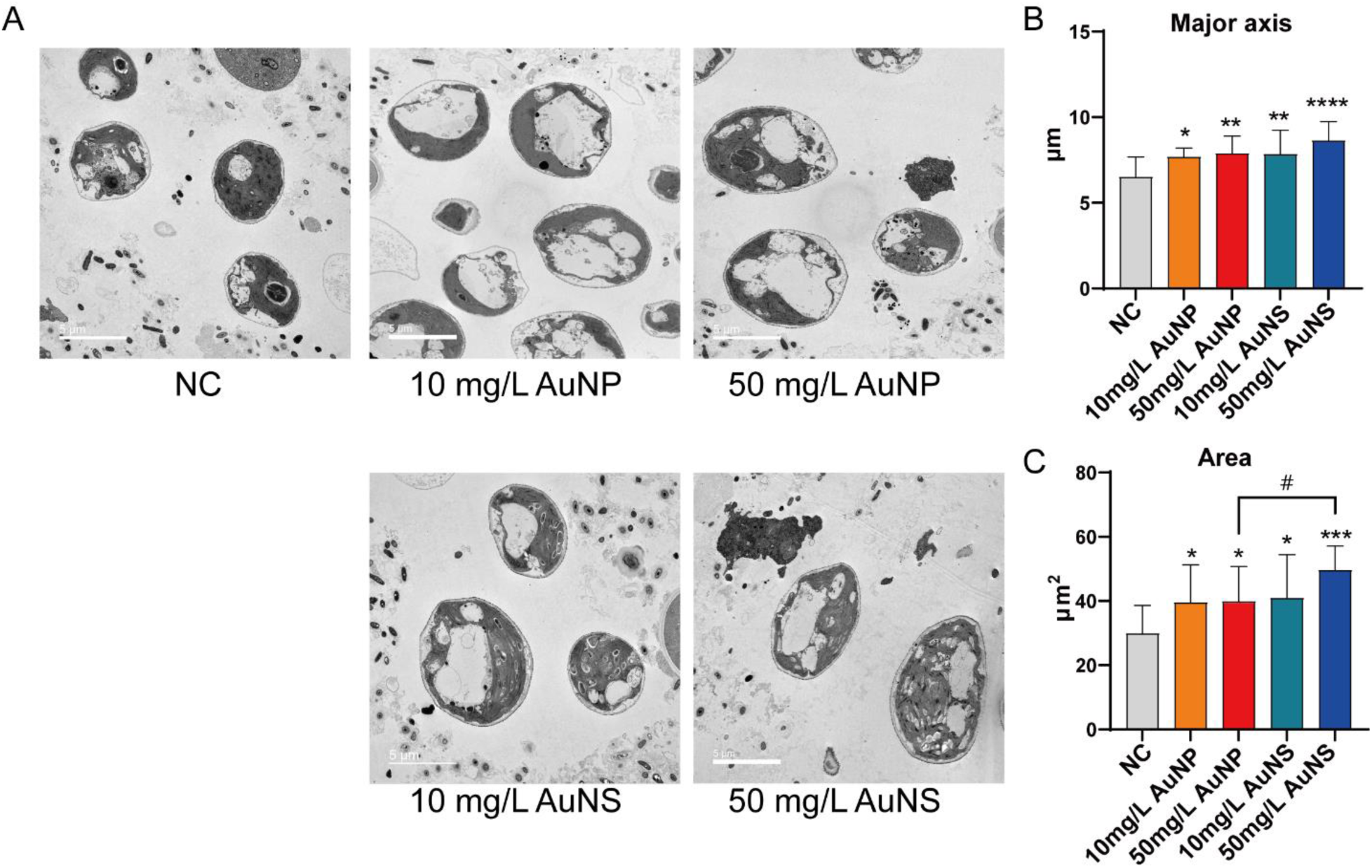
Exposure to AuNP and AuNS enlarged cells size of C. reinhardtii. A. TEM images (2,000X). B. Major axis of single cell. C. Area of single cell. Data are presented as mean ± SD. n = 15. * *p* < 0.05, ** *p* < 0.01, *** *p* < 0.005, and **** *p* < 0.001 compared with control group or between the two groups with lines.

**Figure S4.**
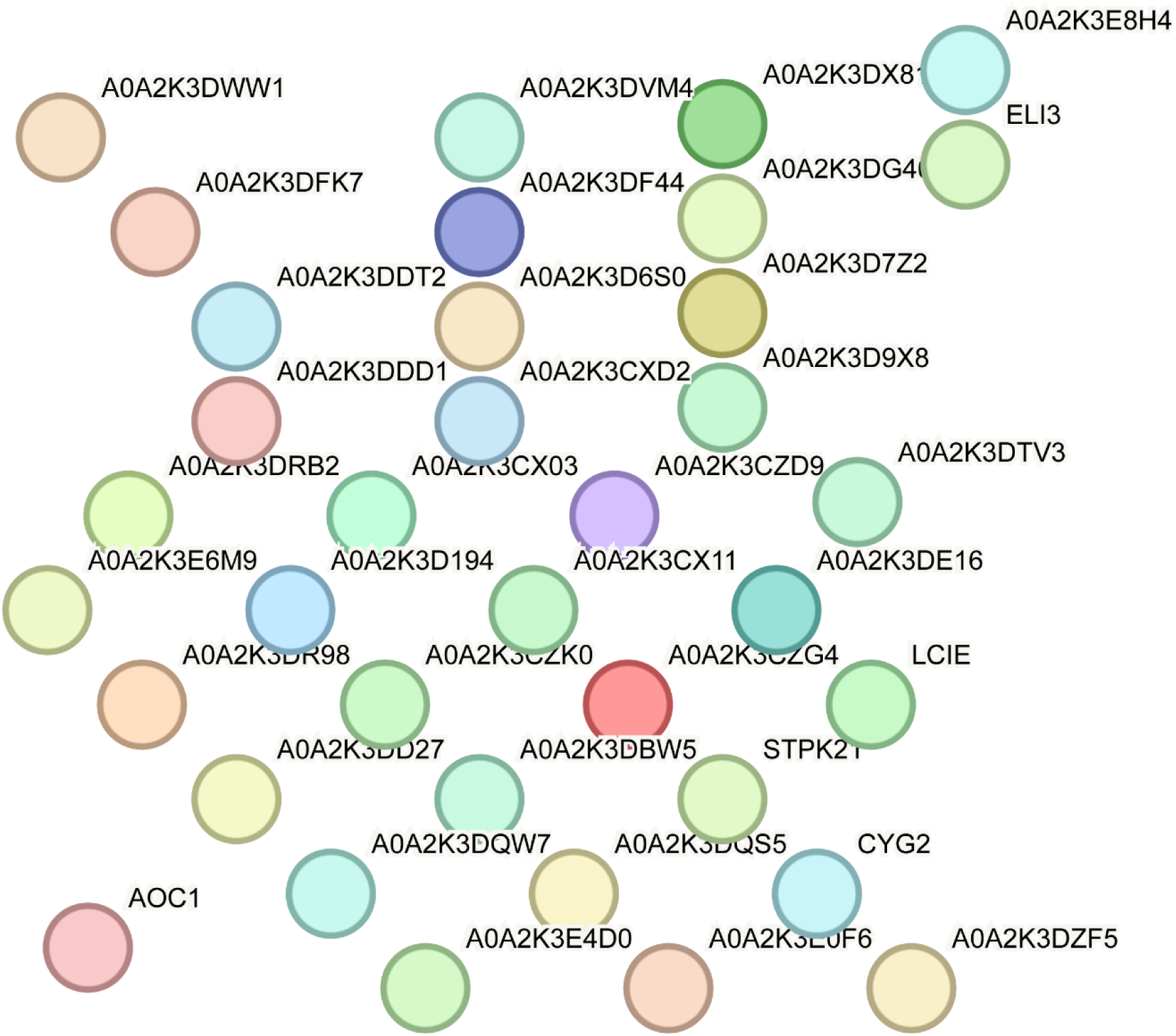
String analysis of common differential expression genes in 10 mg/L AuNP and AuNS groups

